# Smart-design of universally decorated nano-particles for drug delivery applications driven by active transport

**DOI:** 10.1101/2022.06.16.496384

**Authors:** Gal Halbi, Itay Fayer, Dina Aranovich, Ashraf Brik, Rony Granek, Anne Bernheim-Groswasser

## Abstract

Targeting the cell nucleus remains a challenge for drug delivery. Here we present a universal platform for smart design of nano-particles (NPs) decoration that allows recruitment of multiple dynein motors to drive their active motion towards the nucleus. The uniqueness of our approach is based on using: (i) a spacer polymer, commonly Biotin-Polyethylene-glycol-thiol (B-PEG-SH), whose grafting density and molecular weight can be tuned thereby allowing NP transport optimization, and (ii) protein binding peptides, like cell penetrating, NLS, or cancer targeting, peptides. Universal chemistry is employed to link peptides to the PEG free-end. To manifest our platform, we use a SV40T large antigen-originating NLS peptide. Our modular design allows tuning the number of recruited motors, and to replace the NLS by a variety of other localization signal molecules. Our control of the NP decoration scheme, and the modularity of our platform, carries great advantage for nano-carrier design for drug delivery applications.

## Introduction

In the past three decades, the advances of nanotechnology led to the development of a various nano-particles (NP) for drug delivery applications.^1–3^ Commonly, these NPs are based on mesoporous silica materials,^4,5^ lipoplexes,^6^ polyplexes,^7,8^ complexes of peptides, ^9^ magnetic materials,^10–14^ polymer-peptides, ^15–17^ polysaccharides,^18^ protein-DNA complexes,^19–21^ gold nanoparticles,^22–25^ and alginate-sulfate.^26^ Using NPs carries several advantages. First, NPs protect the drug from being attacked and desegregated by the immune and enzymatic systems. Second, due to their nano-size, the NPs can penetrate the pores of the blood vessels and reach distant tissues. Third, using ligation with specific antibodies that recognize and bind to biological markers (receptors), nano-carriers can be designed to target a specific tissue, which result in minimized side effects associated with systematic administration of a therapeutic agents. Fourth, it is possible to enhance cellular uptake of the NPs by manipulating their size and by using ligands, such as cell penetrating peptides (CPPs),^22,27–34^ with main focus on endocytosis. Fifth, some ligands also allow NPs to overcome endosomal escape, which is another important bottleneck for drug delivery applications.

Drug-delivery can be further improved by targeting a specific intra-cellular compartment. By applying such an approach, the drug doses can be reduced, thereby minimizing the disruption of cell function and associated side effects. In contrast, a nano-carrier carried through the cytoplasm *via* diffusion alone is less likely to reach its destination, resulting in significantly larger doses required to elicit the same therapeutic effect. Nevertheless, only few efforts were made to design NPs capable of targeting specific intra-cellular compartments. Among those targets, the nucleus – wherein gene transcription and regulation occurs – shows a great promise for drug delivery applications.^22,31,35–39^ One way of improving NPs’ transport efficiency towards the nucleus is by utilizing the cell’s intrinsic active transport system, in particular, harnessing the microtubule (MT) network and its associated motor protein dynein.^40,41^

Indeed, different types of viruses have evolved to harness the intracellular dynein machinery for efficient targeting of the nucleus, where the infection process of the host cell takes place. HIV, herpes-simplex virus, and Porcine circovirus type 2, express peptide sequences that serve both as nuclear localization signals (NLSs)^42^ and CPPs,^27–30,43–52^ thereby enhancing cellular uptake and transport to the nucleus. An NLS peptide first recruits an *α*-importin protein from the cytosol, followed by binding of a *β*-importin protein, which subsequently recruits a dynein motor.^53,54^ Similarly, adenoviruses use other ligands (e.g., hexon) for engaging the active transport mechanism. ^55–58^ For this reason, these viruses have been successfully used as viral vectors for DNA transfection. One unique example, is the recent use of adenoviruses in various COVID-19 vaccines, in particular the Oxford-AstraZeneca and Johnson & Johnson vaccines, in which the adenovirus envelope is used to deliver the SARS-COV-2 spike protein DNA to the nucleus of muscle cells. A known problem associated with using such viral vectors is the case in which the patient has been previously exposed to the same adenovirus. Under these circumstances, the immune system response prevents transfection of the target cells. Hence, replacing viral vectors with a synthetic nano-carrier with an equivalent nuclear localization is warranted. ^59–64^

In this work, we aspire to mimic the mechanism used by the mentioned above viruses. We aim to design a general NP *decoration* that is independent of the NP material core and can be used for a number of purposes: (i) enhancing the transport efficiency to the nucleus (via NLSs ligation), (ii) enhancing cell penetration (via CPPs ligation), and (iii) tissue targeting, e.g., by using cancer targeting peptides (CTPs).^27–30,51,52,65^ In addition, our decoration is made to provide multiple binding sites for other (yet to be discovered) peptides for targeting, diagnosing, and treating various diseases. We choose in the present study bare NPs made of polystyrene, as they are common and easy to decorate with various ligands, specifically Neutravidin (Ne-Avidin). Clearly, the material from which the bare NP is made of has no influence whatsoever on the properties of the decorated NP, thereby conclusions deduced from this study will be applicable for other bare NP materials.

Our strategy is based on the following decoration steps: (i) Covering the surface of the carboxyl polystyrene NP core by a densely packed Ne-Avidin monolayer, (ii) Binding Biotin-Polyethylene-glycol-thiol (B-PEG-SH) molecules to the Ne-Avidin molecules *via* biotin-avidin bonding (the PEG molecular weight and graft density can be adjusted and optimized for various purposes), and (iii) covalently end-linking a controlled fraction of B-PEG-SHs with a single brominated-modified specialised peptide, e.g., NLS, CPP, CTP etc., or any other desired hybrid (Chimeric) peptide (see Figure 1A).

**Figure 1:**
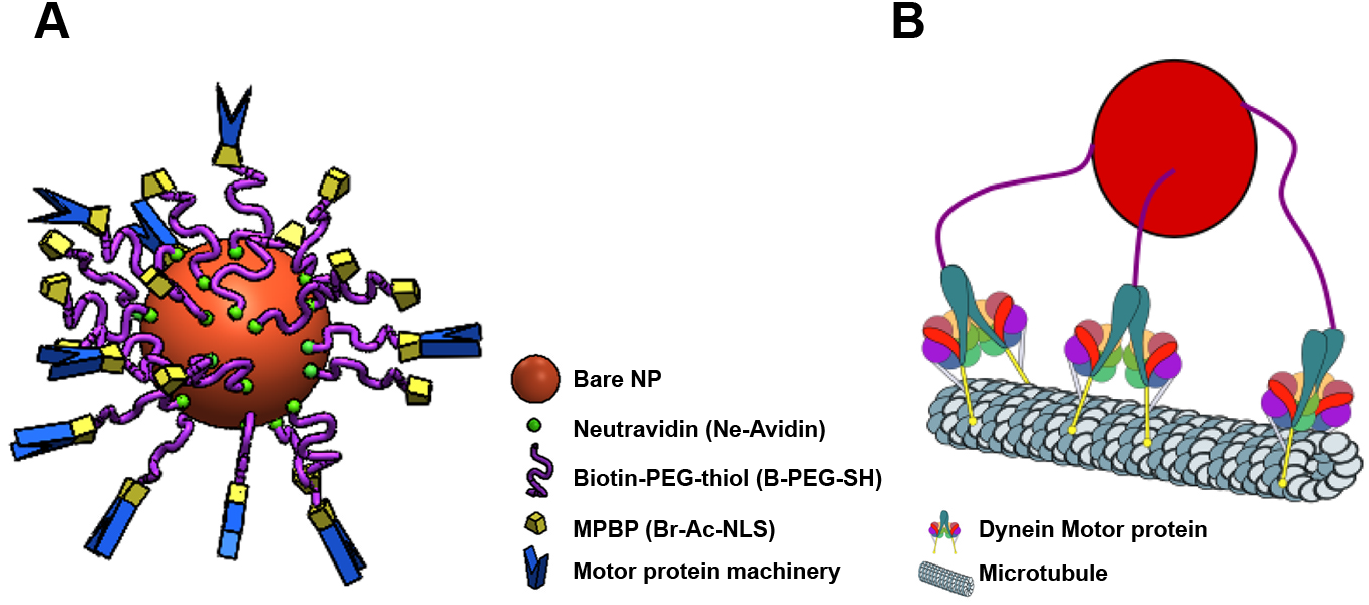
(A) Schematic illustration of the decorated NP (shown a small fraction of decorating molecules). The NP (shown in red) surface is saturated with Ne-Avidin molecules (shown in green). MPBP refers to Motor Protein Binding Peptide. In this work we use a bromo-acetamide (Br-Ac) modified NLS peptide. The motor protein transport machinery includes dynein motor proteins and relevant adaptors (shown in blue). (B) Schematic illustration of a NP having three anchored B-PEG-SH spacer polymers each coupled to a single dynein motor protein. PEG strechability allows the dynein motors to easily interact with the microtubule (MT) curved surface. Color coding as in (A).

Although we demonstrate our strategy only for a particular NLS peptide, the chemistry (SN2 reaction) involved in its binding is identical for all peptides. The B-PEG-SH grafting density, linked NLS fraction, B-PEG-SH contour length, particle size, and concentration of cytoplasmic dynein machinery, are all together expected to govern the number of associated dynein motors that bind to the MT simultaneously (see Figure 1B). As we have shown previously, our designed universally decorated nano-particles, with multiple binding sites for dynein motors, can move actively towards the MT-minus end for distances that are much longer than the single dynein run-length, thereby enhancing the NP transport towards the nucleus.^66^

## Results and discussion

NP synthesis includes several steps. A single component is added at each step (Figure 1A).^66^ The NP core consists of a carboxyl polystyrene (PS-COOH) microsphere of defined diameter. In this work we use both 40 and 200 *nm* diameter microspheres. The total surface area of the microspheres is kept constant in all experiments and is the same for all microspheres, regardless of their diameter. Via this, we guarantee that the surface density of all added components, notably, PEG-NLS molecules, is independent of the NP core size.

First, the bare NPs (i.e., microsphere) are incubated with 10 *µM* Ne-Avidin to saturate the NP surface with Ne-Avidin molecules. ^67^ The positively charged Ne-Avidin molecules bind to the negatively charged (polystyrene) NP surface *via* physical electrostatic interactions. The NPs are then passivated with Bovine Serum Albumin (BSA) to block possible non-specific adsorption of molecules in subsequent decoration steps. The NPs are then incubated with 1*mM* B-PEG-SH, 100-fold higher than Ne-Avidin. Moreover, since the Ne-Avidin-decorated NPs attach to the B-PEG-SH molecules *via* biotin-avidin bonding, the strongest known non-covalent interaction between a protein and its ligand (*K*_*d*_ = 10^−15^M) we expect that each B-PEG-SH binds to a Ne-Avidin molecule such that a protective PEG shell is formed on the NP surface. This protective layer plays a critical role in preventing aggregation of adjacent NPs and in enhancing their stability in the route towards the targeted organ/cell. The PEG decorated NPs are then incubated with a solution of bromo-acetamide modified (Br-Ac) SV40T large antigen NLS peptides. The Br-Ac group is followed by a GGGG amino acid sequence (raft) that spatially separate the reactive Br-Ac group from the NLS peptide (Figure 2A). The thiol group of the B-PEG-SH molecule react with the Br via a SN2 reaction which covalently attaches the NLS to the NP surface (Figure 2A). Noteworthy, this reaction can be used to bind other peptides under a permissive range of conditions such as pH, temperatures, and solution ionic strength. The raft can also functions as a regioselective addressable functional template that allows the incorporation of functional units, e.g., a fluorescent dye. In this case, we use a slightly modified raft which includes a Lys (K) side chain (GGGKGGG) to which 5-carboxytetramethylrhodamine (TAMRA) is linked^68^ to obtain GGGK(TAMRA)-GGGNLS (TAMRA-NLS) (Figure 2B).

**Figure 2:**
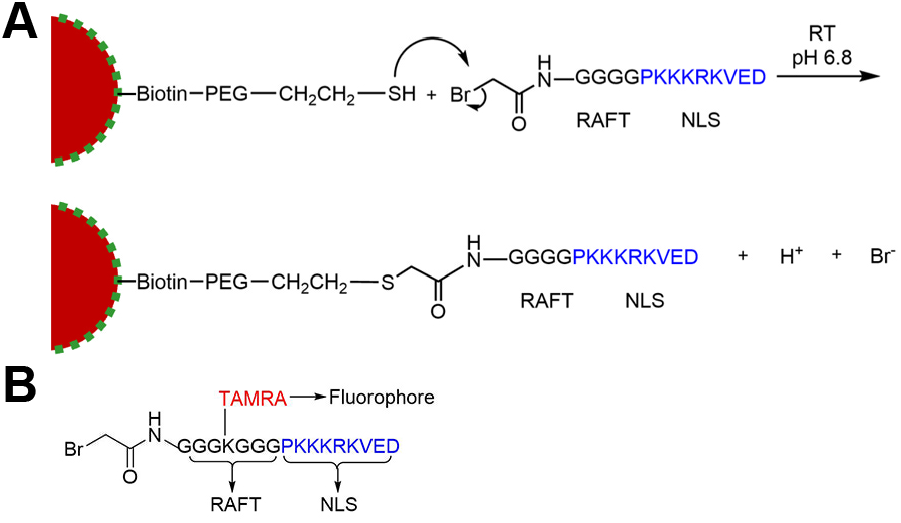
(A) Covalent attachment of the NLS peptide to the NP surface occurs *via* a SN2 reaction between Br of the bromo-acetamide (Br-Ac) modified NLS and the thiol group of the B-PEG-SH molecule. Color coding as in Fig. 1A. (B) Sequence and chemical formula of the fluorescently labeled NLS peptide (TAMRA-NLS).

To determine the effect of each added component on NP decoration, the NPs are characterized using different techniques. First, we study the effect of B-PEG-SH molecular weight, *M*_*w*_, on the properties of the protective shell formed on the NP surface by measuring the inter-particle distance between adjacent B-PEG-SH decorated NPs of increasing *M*_*w*_ using cryo-TEM (Figure 3A; see (d) for illustration). The black dots on the particle’s surface are the Ne-Avidin molecules, whose diameter, *D*_*Ne*−*Avidin*_, is about 5 *nm*.^69^ NPs not grafted with B-PEG-SHs are used as a control group (Figure 3A(a)). In the absence of B-PEG-SH spacer polymers, the mean spacing between adjacent NPs is 10.5 ± 7 *nm* (mean ± STD) (Figure 3B), typically twice the diameter of a Ne-Avidin molecule.^69^ The inter-particle distance increases as a function of the B-PEG-SHs *M*_*w*_ and reaches saturation for B-PEG-SH of 5 *kDa*, for which the distance between adjacent NPs is 26 ± 5 *nm* (mean ± STD). This distance is significantly larger than the characteristic gyration radius of a single, free PEG distance is significantly larger than the c polymer gyration radius, 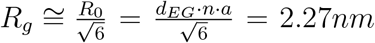, where *R*_0_ is the free polymer end-to-end distance,^70,71^ *d*_*EG*_ = 0.358 *nm* is the ethylene glycol subunit length, *a* = 0.76 *nm* is the PEG Kuhn length,^70,72^ and *n* = 114 is the number of ethylene glycol subunit for a 5 *kDa* B-PEG-SH. Since the Ne-Avidin molecules are closely packed on the NP surface, we conclude that the anchored B-PEG-SH molecules are forming a protective layer on the NP surface of densely packed random coils (sometimes referred to as “mushrooms”).^73^ All subsequent experiments are performed with 5 *kDa* B-PEG-SH.

**Figure 3:**
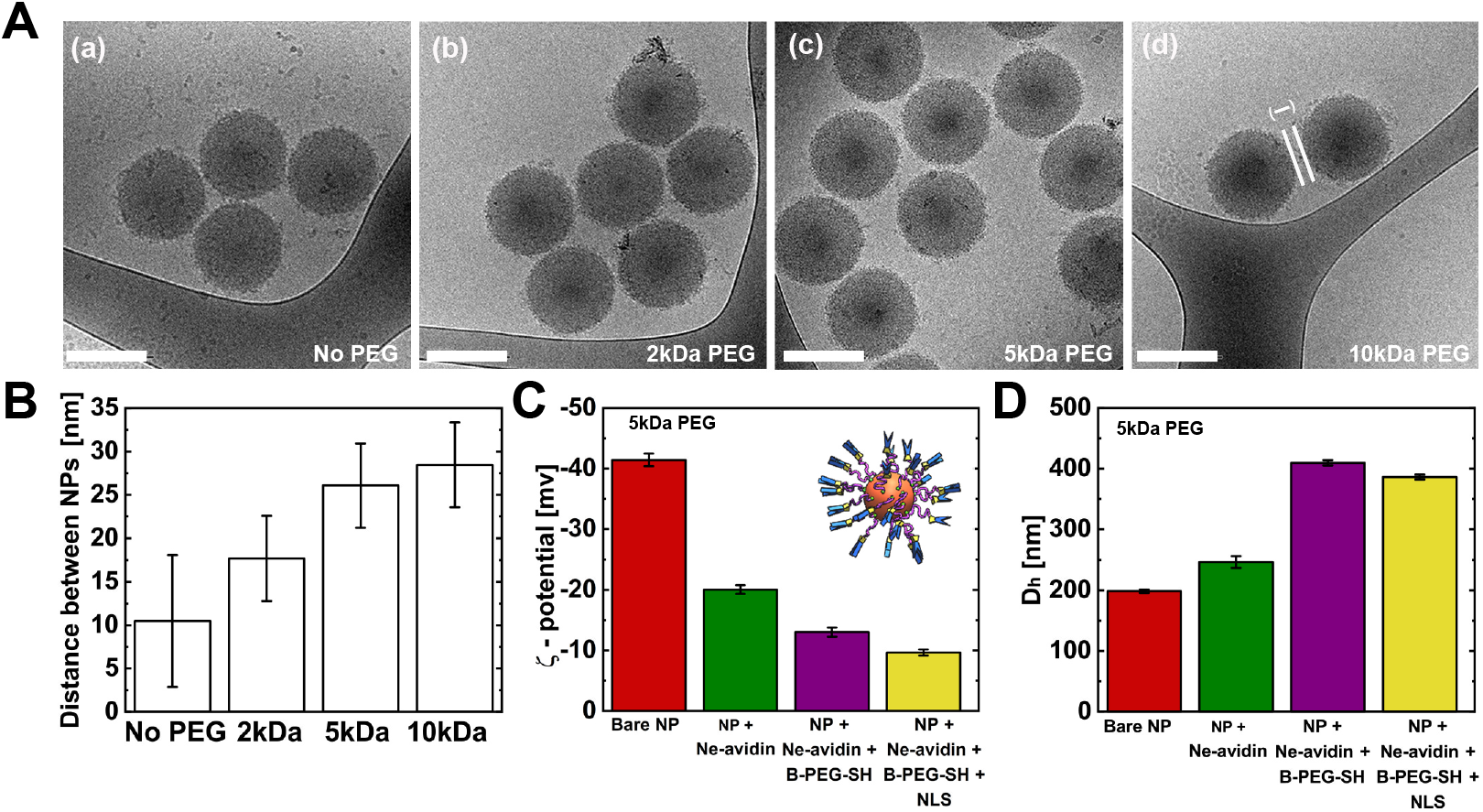
NP characterization. (A) Cryo-TEM images of NPs decorated with Ne-Avidin molecules (a) and NPs decorated with Ne-Avidin and with B-PEG-SH of increasing molecular weights: *M*_*w*_ is 2 *kDa* (b), 5 *kDa* (c), and 10 *kDa* (d). Scale bars are 200 *nm*. (B) Mean distance between adjacent NPs for B-PEG-SH of different *M*_*w*_. The distance is determined as shown in panel A(d). (C) *ζ*-potential values and (D) hydrodynamic diameter, *D*_*h*_, as a function of added components (column) with same color coding as in Fig. 1A. See also schematic NP in the inset of (C). Values in (B-D) correspond to mean ± STD. (A-D) NPs have a mean core diameter of 200 *nm*.

To ensure the correct integration of the different decoration components, we utilize *ζ*-potential and DLS assays to examine how the NPs charge and effective diameter change throughout the assembly steps, respectively. Zeta potential measurements show that the bare NP microspheres are highly negatively charged (*ζ*-potential value of −41.4 ± 1 *mV*, mean ± STD) (Figure 3C). While the *ζ*-potential values remain negative along the various decoration stages, they gradually decrease in absolute values. This trend is consistent with the assumption that both Ne-Avidin and NLS molecules are positively charged (see Materials and methods); thus, a reduction of the absolute value of the *ζ*-potential is expected upon their binding.

The DLS data (see Figure 3D) shows a significant increase of the NP hydrodynamic diameter (*D*_*h*_) upon addition of Ne-Avidin and B-PEG-SH (5 *kDa*) compared to the bare NPs. Upon NLS binding, a slight decrease of the *D*_*h*_ is detected, confirming B-PEG-SH binding to the NP surface. These results are consistent with our expectation that the NP hydrodynamic diameter should increase as NP decoration progresses. The decrease of the hydrodynamic diameter upon NLS addition is probably due to the attractive electrostatic interactions between the overall negatively charged NP surface and the positively charged NLS, leading to a small compression of the mushroom-like PEG-NLS complex.

To quantify the amount of attached PEG-NLSs per NP, ⟨*N* ⟩, and the mean anchoring distance between adjacent PEG-NLS molecules, *ξ*^∗^, we carry out UV-Vis absorption experiments using TAMRA-NLS. We incubate the B-PEG-SH grafted NPs with increasing amounts of TAMRA-NLS and measure the absorption of the remaining TAMRA-NLS molecules that *do not* adsorb to the NPs (i.e., supernatant), from which we deduce the total number of TAMRA-NLS molecules that *do* adsorb to the NPs surface, Σ*N*. The mean number of bound PEG-NLSs per NP is then calculated from

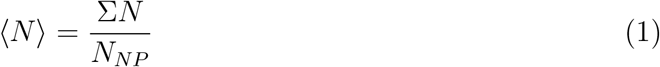

where *N*_*NP*_ is the total number of NPs. For the two NP diameters tested, the dependence of ⟨*N* ⟩ on [TAMRA-NLS] follows a Langmuir-like isotherm (Figure 4A). The mean anchoring distance between adjacent PEG-NLS molecules, *ξ*^∗^, is determined assuming that each PEG-NLS molecule occupies a square lattice site *ξ*^∗2^ on the NP surface

**Figure 4:**
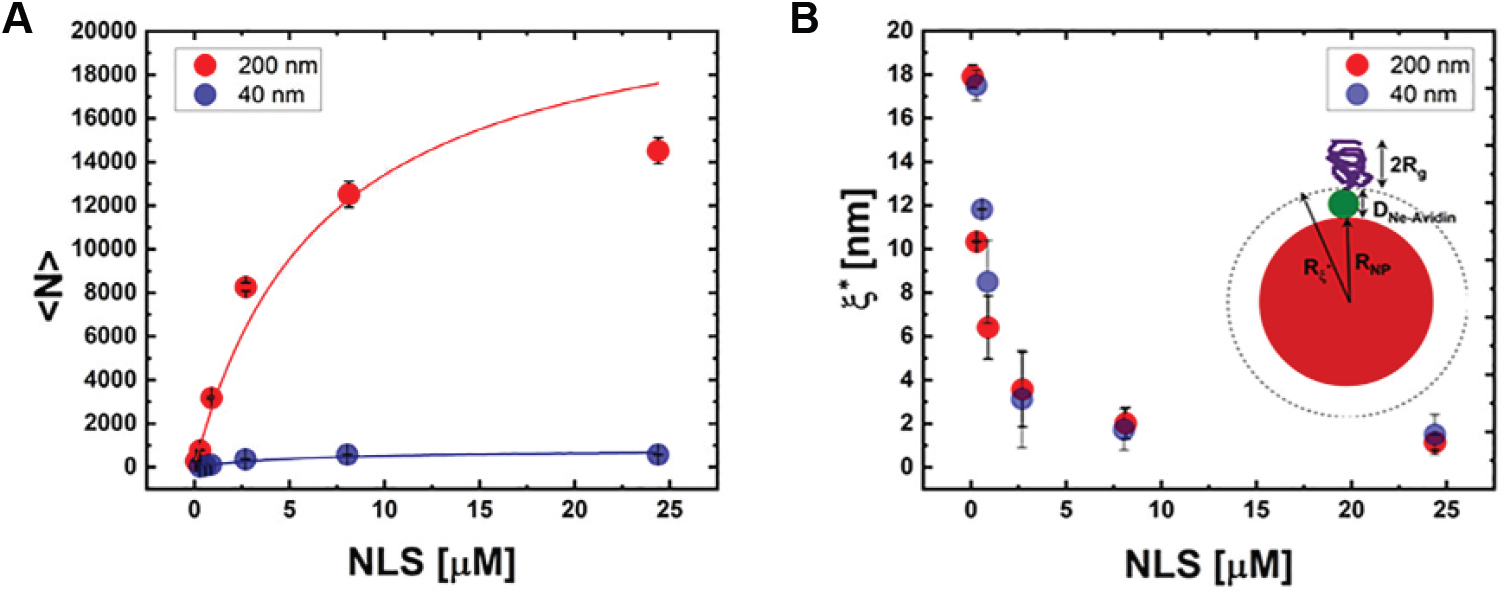
Mean number of grafted PEG-NLSs and mean anchoring distance. (A) Mean number of grafted PEG-NLSs, *N*, against TAMRA-NLS concentration (marked as NLS). The experimental data (dots) follow a Langmuir-like isotherm (line) with *R*^2^ = 0.95 and 0.96 for the 40 and 200 *nm* core diameter NPs, respectively. (B) Mean anchoring distance between neighboring PEG-NLSs, *ξ*^∗^, determined at the surface of a sphere of effective radius *R*_*ξ*_*∗* (inset), versus [TAMRA-NLS]. (Inset) schematic illustration of the NP with the relevant characteristic dimensions: *R*_*NP*_ - NP core radius, *D*_*Ne*−*Avidin*_ - Ne-Avidin diameter, and *R*_*g*_ - PEG gyration radius. Color coding as in Fig. 1A.

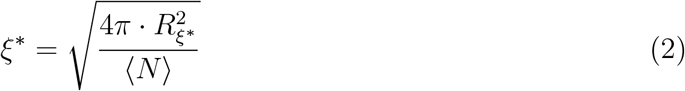

where *R*_*ξ*_*∗* is the mean radius of Ne-Avidin coated NPs (see inset in Figure 4B). *ξ*^∗^ gradually decreases with the increase in [TAMRA-NLS] and appears to be independent of the NP core diameter, see Figure 4B. *ξ*^∗^ reaches a minimal finite value of 4.1 ± 0.7 *nm* at 4 *µM* TAMRA-NLS, of the same order of magnitude as the PEG diameter, 2 · *R*_*g*_ = 4.5*nm*. This suggests that at saturating conditions each B-PEG-SH molecule has a NLS molecule attached to it. Note that the observed independence of *ξ*^∗^ on the NP core diameter confirms the scalability of NP decoration.

Having confirmed the proper integration of the various added components, we now turn to quantify the amount of recruited dynein motors per NP. Recruitment of the *mammalian* dynein motor proteins machinery, which is mediated by NLS,^66^ is achieved by incubating the PEG-NLS-coated NPs in Hela cytoplasmic cells extract (CE). ^74,75^ The amount of recruited dynein motors, which we expect to depend on the number of bound NLSs, and thus on [NLS], is quantified using WB assays. We prepare samples in a range of [NLS] = 0.1-2.7 *µM*, in which the most significant changes in the number of attached NLSs was detected (Figure 4A). Exposure of the 200 *nm* NPs to CE demonstrates antibody-specific binding to *mammalian* dynein motors (Figure 5A). We quantify the total amount of recruited dynein motors, Σ*N*_*m*_, from the integrated signal measured from the area of the dynein bands, which we calibrate using purified dynein proteins (see Methods), from which we then deduce the mean number of bound dynein motors per NP

**Figure 5:**
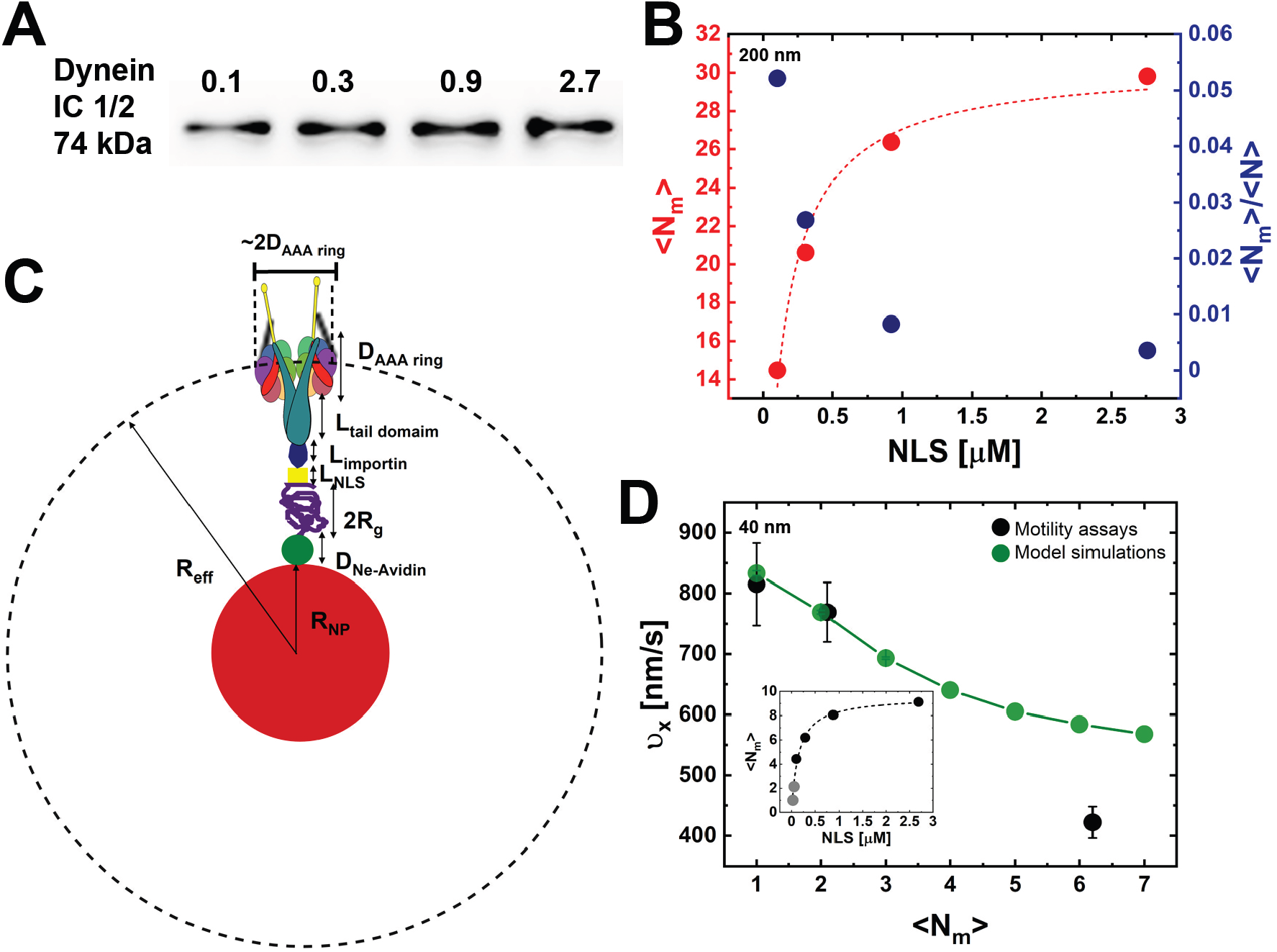
Quantifying the amount of recruited *mammalian* dynein motors. (A) Western blot bands demonstrating *mammalian* dynein recruitment to the NPs surface against [NLS]. (B) Mean number of recruited *mammalian* dynein motors per NP, ⟨*N*_*m*_⟩, versus [NLS]. The experimental data (red dots) follows a Langmuir-like isotherm (red dotted line) with *R*^2^ = 0.98. Fraction of NLSs occupied by dynein motor proteins, <u>^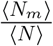^</u>, versus [NLS] (blue dots). (A,B) Mean NPs core diameter is 200 *nm*. (C) ⟨*N*_*m*_⟩ is evaluated at the surface localized at *R*_*eff*_, as illustrated in the figure. Specifically *R*_*eff*_ = *R*_*NP*_ + *D*_*Ne*−*Avidin*_ + 2 · *R*_*g*_ + *L*_*NLS*_ + *L*_*importin*_ + *L*_*tail domain*_ + 0.5 *D*_*AAA ring*_ where *R*_*NP*_, *D*_*Ne*−*Avidin*_, and *R*_*g*_ are as in Fig. 4B (see main text for actual dimensions); *L*_*importin*_ = 15 *nm* - length of the *αβ*-importins complex, ^54^ *L*_*tail domain*_ = 39 *nm* - length of the relevant part of the dynein tail domain,^79^ *D*_*AAA ring*_ = 13 *nm* - dynein AAA ring diameter,^80^ and *L*_*NLS*_ - length of the NLS peptide, which is neglected. Color coding as in Fig. 1A. (D) Black symbols: Mean longitudinal velocity extracted from bead motility assays^66^ against the mean number of recruited *mammalian* dynein motors estimated using the data presented in the inset and the data shown in Fig. 4A in Ref.^66^ Green symbols and connecting line: model simulations results for the mean longitudinal velocity against the number of NP-bound motors taken from Ref. ^66^ (Inset) Mean number of recruited *mammalian* dynein motors per NP ⟨*N*_*m*_⟩ versus [NLS]. The experimental data (dots) follows a Langmuir-like isotherm (line) with *R*^2^ = 0.99. The two bright grey dots correspond to extrapolated values calculated from the fit, for [NLS] = 0.025 and 0.05 *µM*. Mean NPs core diameter is 40 *nm*.

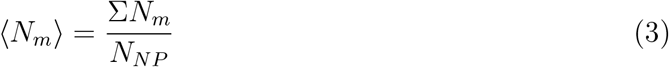

where *N*_*NP*_ is the total number of NPs. The dependence of ⟨*N*_*m*_⟩ on [NLS] follows a Langmuir-like isotherm similarly to ⟨*N* ⟩, yet, saturation is reached at slightly lower [NLS] = 2 *µM* (Figure 5B). We also notice that only a fraction of the attached NLSs, 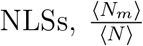, are occupied by dynein motor proteins, suggesting that the number of attached NLSs represents an upper bound for the amount of attached dynein motors (Figure 5B). Note that this fraction further decreases as the amount of attached NLSs increases. We can reasonably assume that since the surface density of the PEG-NLSs, *ξ*^∗−2^, is similar for NPs of different diameters, the surface density of the recruited dynein motors should be also the same. Based on this, we can estimate the mean number of dynein motors recruited to the surface of the 40 *nm* diameter NPs, ⟨*N*_*m*_⟩_40*nm*_ (Figure 5D, inset), from the mean number of recruited dynein motors on the 200 *nm* diameter NPs, ⟨*N*_*m*_⟩_200*nm*_, taking also into account the difference in the NPs surface areas:

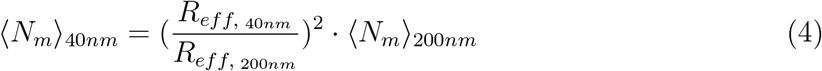

The relevant surface areas are evaluated at *R*_*eff*_, which defines the radius where the dynein motor complexes occupied area is the widest such that the repulsive excluded volume interactions between neighboring motor complexes is the largest, see Figure 5C, which is expected to limit the number of motor complexes that can accommodate on the NP surface. We can now turn to ask how the mean longitudinal velocity of the 40 *nm* NPs, ⟨*v*_*x*_⟩, extracted from bead motility assays^66^ depends on the number of recruited motors, see Figure 5D. Our data show that at [NLS] = 0.025, 0.05, and 0.3 *µM*, the NPs are moving with a mean number of recruited motors of ⟨*N*_*m*_⟩ = 1.8, 3.2, and 6.2, respectively (Figure 5D, black dot); this is in good agreement with our predicted values for the mean velocity against the number of NP-bound motors obtained from Monte-Carlo (MC) simulations (Figure 5D, green dot).^66^

## Conclusions

In this paper, we have presented a general platform for the smart design of nano-particle decoration for nuclear drug delivery applications. The most essential parts of the design scheme are: (i) the incorporation of spacer polymers, commonly PEG, and (ii) the use of a protein binding peptide, such as CPP, NLS peptide, or CTP, to assist for the variety of cellular processes involved in the drug delivery. The B-PEG-SH grafting density over NP surface and its molecular weight can be accurately and readily controlled. We posit that adjusting these parameters will influence the associated cellular drug delivery assisting processes, thereby allowing for their optimization. Furthermore, the chemistry that links the peptides to the PEG free-end is not exclusive for the given peptide and is easy to apply, even for a mixture of peptides (e.g. some PEGs will end with an NLS, some with a CPP, and others with a CTP), resulting in a flexible design.

To manifest our platform, we have chosen a SV40T large antigen-originating NLS peptide which can recruit dynein motor proteins from the cytoplasm, as well as facilitating the NP penetration through the nuclear pore. ^76–78^ We presented an elaborate characterization of all steps of synthesis. As dynein walks from the cell periphery towards the centrosome (where the MTs minus-ends concentrate), which is located at the vicinity of the nucleus, the NLS is expected to enhance nuclear co-localization. Since many PEG-NLSs cover the NP surface, even if just a fraction of them recruit dynein, the intracellular run-length and run-time for the NP motion over an MT are expected to be greatly improved, as we have showen recently. ^66^ Therefore, if the NP core will be made from a bio-compatible drug carrier material (instead of the polystyrene used here), it is high-likely that the drug delivery to the nucleus will be amplified.

Further studies of the sensitivity of motility properties on the PEG *M*_*w*_ and grafting density will allow further optimization of nuclear drug delivery. Similar optimization studies, associated with PEG *M*_*w*_ and grafting density variations, can be performed for the endocytosis processes when CPPs are used, or for cancer cell targeting when using CTPs. In doing so, we believe that our control of the specific NP decoration studied here, and our (general) controlled platform proposition, carry great promise for nano-carrier design for drug delivery applications. Furthermore, our NP modular design allows us to easily tune the NPs transport motility properties by controlling the number of motor proteins recruited to the NP surface. Finally, one can replace the NLSs with a variety of other localization signal molecules and motor protein binding peptides, thereby opening the way to a wide range of medical applications.

## Acknowledgement

This research was initially supported by the Focal Technological Area Program of the Israeli National Nanotechnology Initiative (INNI). The authors thank Dr. Einat Roth Nativ and Dr. Alexander Upcher for NP imaging by cryo-TEM.

## Supporting Information Available Materials and Methods

### Materials

#### Hela cell extracts preparation

Hela cell extracts were prepared according to Fu et al. ^81^ Briefly, Hela cells were grown in DMEM medium supplemented with 10 % of Fetal Bovine Serum (FBS) and 1 % of penicillin-streptomycin at 37 °C and 5 % CO_2_. The cells were detached by Trypsin-EDTA solution, washed with Phosphate Buffered Saline (PBS) (10 *mM* Phosphate buffer, 2.7 *mM* KCl, 137 *mM* NaCl, pH = 7.4), and pelleted for 5 *min* at RT and 500 g. The cells were then incubated on ice with a lysis buffer (12 *mM* Pipes, 2 *mM* MgCl_2_, 1 *mM* EGTA, 0.4 % Triton X-100, 10 % protease inhibitor cocktail, pH = 6.8) for 15 *min*. Finally, the lysates were cleared by centrifugation at 2,700 *g* and then at 100,000 *g*, at 4 °C for 10 *min* each. The cell extracts were diluted 10x with lysis buffer without Triton and protease inhibitors. The total protein mass concentration was determined by Bradford. Sucrose was added to the extracts (10 % in mass), flash-frozen in liquid nitrogen and stored at −80 °C.

#### Nuclear Localisation Signal (NLS) peptide

The NLS sequence (PKKKRKVED) originates from SV40T large antigen.^82,83^ We use a N-terminal bromo-acetamide (Br-Ac) modified NLS prepared using solid phase peptide synthesis^68^ which under our working conditions (pH = 6.8-7.0) is positively charged. The bromine-acetamide group is followed by an amino acid sequence that functions as a regioselective addressable functional template (RAFT) that allows the incorporation of functional units. We use a GGGKGGG raft to label the NLS fluorescently. A shorter raft (GGGG) is used for the non-labeled NLS. Specifically, 5-carboxytetramethylrhodamine (TAMRA) is linked to the Lys side chain^68^ to obtain GGGK(TAMRA)-GGGNLS (TAMRA-NLS), which is purified by HPLC and characterized by mass spectrometry. The absorbance maxima of TAMRA-NLS is at 553 *nm*.

#### NPs preparation

Fluorescent Carboxyl Polystyrene (PS-COOH) Microspheres of 40 and 200 *nm* in diameter (FCDG001 and FCDG003, Bangs Laboratories, Fishers, IN, USA) are used for NPs preparation. The total surface area of the microspheres is kept constant in all experiments, and is same for all microspheres regardless of their diameter. The various incubation steps are performed at 25 °C for 20 *min* in 1 *mM* Hepes (pH 6.8) and a total volume of 50 *µL*, unless stated otherwise. Specifically, 2 *µL* of 40 *nm* diameter microspheres (or 10 *µL* of 200 *nm* diameter microspheres) are incubated with 10 *µM* Ne-Avidin (31000, Thermo Fisher). Ne-Avidin, which is positively charged, strongly binds to the negatively charged (polystyrene) NP surface *via* physical electrostatic interactions. Excess Ne-Avidin is separated by 3 dialysis cycles. Next, the NPs are incubated with 10 *mg/mL* Bovine Serum Albumin (BSA) to block non-specific adsorption in subsequent decoration steps. Excess BSA is separated by centrifugation (6800 *g* for 8 *min*). All subsequent centrifugation steps are performed at the same conditions. The NPs are then incubated with 1 *mM* B-PEG-SH (NANOCS, New York, NY, USA) for 30 *min*. Excess B-PEG-SH is separated by centrifugation. We covalently attach the NLSs to the NPs surface by incubating the NPs in a solution of Br-Ac modified NLS in PBS. Excess NLSs are separated by centrifugation. In the experiments we vary the concentration of NLS between 0.1-25 *µM*. Finally, the NPs are incubated with 20 *µL* cytoplasmic Hela cell extracts at 30 °C for 20 *min* allowing the recruitment of the *mammalian* dynein motor machinery to the NPs surface. The NPs are washed twice and kept on ice in 20 *µ*L BRB80 solution (80 *mM* PIPES pH 6.8, 1 *mM* MgCl2,1 *mM* EGTA) supplemented with 1 *mM* Mg-ATP.

## Methods

### Cryo-TEM

Cryo-TEM samples are produced using a standard process. ^84^ Shortly, vitrified specimens are prepared by applying 2.5 *µL* drop of the NPs solution on a copper grid coated with a perforated lacy carbon 300 mesh (Ted Pella Inc, Canada, USA), then blotted with a filter paper to form a thin liquid film. The grid is immediately plunged into liquid ethane at its freezing point (−183 °C) using an automatic plunger (Leica, EM GP, Buffalo Grove, IL, USA). The vitrified specimens are analyzed using a FEI Tecnai 12 G2 TEM at 120 *kV* with a Gatan cryo-holder maintained at −180 °C. Images are recorded on a CCD camera at low dose conditions and recording is performed using Digital Micrograph software (Gatan manufacturer, Pleasanton, CA, USA).

### DLS and *ζ*-potential experiments

DLS and *ζ*-potential measurements are performed on a Malvern NanoZS instrument (ZN-NanoSizer, Malvern, England) operating with a 2 *mW* HeNe laser at a wavelength of 632.8 *nm*. The detection angle is 173 ° and 17 ° for DLS and *ζ*-potential measurements, respectively. All measurements are done in a temperature-controlled chamber at 25 °C (± 0.05 °C); for data analysis the viscosity is taken as that of water (0.8872 *cP*). The NPs hydrodynamic diameter *D*_*h*_ is extracted as follows: the intensity size distribution is extracted from the intensity auto-correlation function calculated using an ALV/LSE 5003 correlator over a time window of 30 sec (10 runs of 3 sec), repeated 3 times, using the software CONTIN. For *ζ*-potential measurements, the solution is transferred to a U-tube cuvette (DTS1070, Malvern, England); the instrument is operated in automatic mode. Utilizing the measured NPs electrostatic mobility, the *ζ*-potential value is determined by applying the Henry equation. ^85^ The *D*_*h*_ and *ζ*-potential values are averaged over 3 independent experiments; error bars correspond to the standard deviations of experimental values.

### UVVis absorption experiments

UVVis absorption experiments are used to determine the mean number of bound PEG-NLS molecules per NP, ⟨*N* ⟩. Analysis of the data is done using an extinction coefficient that we extract from the absorbance of solutions of increasing TAMRA-NLS concentrations that we fit using Beer-Lambert law. The absorbance is measured at a wavelength of *λ* = 553 *nm* resulting in an extinction coefficient *ϵ*_553_ = 0.0639 [1/(*µMcm*)].

### Western blot experiments

WB is used to confirm the recruitment of dynein motors by the PEG-NLS coated NPs and to quantify the amount of *mammalian* dynein molecules recruited to the NP surface. The NPs are prepared as detailed above, then pelleted at 6800 *g* for 10 *min* at 25 °C, resuspended in a 1× Laemmli Sample Buffer, and boiled for 5 *min* to promote the detachment of the bound proteins. The proteins are separated by electrophoresis using a 12 % agarose gel and transferred to a nitrocellulose membrane. The membrane is incubated for 1 *h* in a blocking buffer of PBST (PBS supplemented with 0.1 *v/v* % Tween) and 10% (*v/w*) dry skim milk (Sigma-Aldrich, St. Louis, MO, USA). The membrane is washed three times with PBST for 5 *min* and then incubated for 1 *h* at 25 °C with a primary anti-dynein antibody (sc-13524) (Santa Cruz Biotechnology, Dallas, TX, USA) diluted 1:200 (*v/v*) in blocking buffer prior use. Next, the membrane is washed three times with PBST and incubated for 1 h at 25 °C with an anti-mouse HRP conjugated with a secondary antibody (sc-2005, Santa Cruz Biotechnology, Dallas, TX, USA) diluted 1:10,000 (*v/v*) in PBST supplemented with 0.5% (*v/w*) skim milk. To finalize the procedure, the membrane is washed three times with PBST and incubated with an ECL Western blot reagent (1,705,060, Bio-Rad, Hercules, CA, USA) for 5 *min* in the dark. Images are collected by chemiluminescence using a Fusion FX imaging system (Vilber Lourmat, Collégien, France). The integrated signal measured from the area of the dynein bands is extracted using ImageJ and calibrated using known masses of purified dynein protein (CS-DN01, Cytoskeleton, Denver, CO, USA) that serve as standards. A molecular weight of 2.17 *MDa*^86^ is then used to derive the number of dynein motors recruited to the NPs. The standards and NP samples are prepared under identical conditions.

